# Analyzing Dwell Times with the Generalized Method of Moments

**DOI:** 10.1101/318717

**Authors:** Sadie Piatt, Allen C. Price

## Abstract

The Generalized Method of Moments (GMM) is a statistical method for the analysis of samples from random processes. First developed for the analysis of econometric data, the method is here formulated to extract hidden kinetic parameters from measurements of single molecule dwell times. Our method is based on the analysis of cumulants of the measured dwell times. We develop a general form of an objective function whose minimization can return estimates of decay parameters for any number of intermediates directly from the data. We test the performance of our technique using both simulated and experimental data. We also compare the performance of our method to nonlinear least-squares minimization (NL-LSQM), a commonly-used technique for analysis of single molecule dwell times. Our findings indicate that the GMM performs comparably to NL-LSQM over most of the parameter range we explore. It offers some benefits compared with NL-LSQM in that it does not require binning, exhibits slightly lower bias and variance with small sample sizes (N<20), and is somewhat superior in identifying fast decay times with these same low count data sets. Our results show that the GMM can be a useful tool and complements standard approaches to analysis of single molecule dwell times.

## Introduction

A fundamental challenge in analysis of single molecule dwell times is the determination of the correct model to explain observations. Methods that have been developed to address this fundamental problem include Bayesian inference (1) and maximum likelihood (2), as well as the more common nonlinear least-squares minimization approach (3). Typically, models depend on unknown parameters which are not experimentally accessible. Methods are distinguished by the manner in which these hidden parameters are estimated. In this work, we develop and characterize the use of the Generalized Method of Moments. This method is a statistical estimation method originally developed in econometrics (4).

Single molecule methods generate high resolution quantitative data on biomolecular mechanisms. The advantages of single molecule experiments include identification of rare intermediates and solution of the “dephasing” problem (5). In techniques such as single molecule fluorescence, a change in signal level can indicate a state transition. Other methods, such as particle tracking using micro-beads, can also yield data on single molecule state transitions (6). In general, the determination of how many states are present and how long the molecule dwells in each state can be challenging, and several methods have been developed to deal with this problem (7, 8). Here, we do not treat this problem, but address the problem of the statistical analysis of the dwell times independent of the method in which they are determined. Analysis of the dwell times is a key step in data interpretation and yields statistics on molecular transitions, thus providing data to test competing mechanistic models. The most common method of analysis, nonlinear least-squares minimization (NL-LSQM), is known to produce biased and non-normally-distributed estimates of the model parameters (3).

The Generalized Method of Moments (GMM) is a powerful statistical method for analysis of samples of random processes. From its original application in the modeling of capital asset pricing (4), its use has grown to make it one of the central methods of econometric analysis (9). In spite of its broad applicability, its use in the natural sciences has been quite limited, although it has been applied recently to the analysis of simulated stochastic chemical reaction networks (10). In the GMM, the *population* moments of a random variable, as calculated from a suitable model, are compared to the *sample* moments as calculated from the measured data. Estimates of hidden parameters are determined via minimization of an objective function which quantifies the disagreement between the population and sample moments (9). The GMM is a general framework that can accommodate a wide range of models and data types. Furthermore, in the limit of a large number of samples, the GMM produces normally-distributed and unbiased estimates (4). In the method, multiple moments can be considered, each moment resulting in a constraint on the parameters of the model. In the case of under-constrained systems (number of moments less than number of parameters), non-unique solutions exist. When the number of moments is equal to the number of parameters, a unique set of estimates for the parameters can be determined (assuming a real solution exists). This exactly-determined system is often referred to as the Classical Method of Moments (CMM). In the case of an over-constrained system (number of moments greater than the number of parameters), only an approximate solution can be found in general. The GMM is a framework in which all such cases can be handled.

In this work, we develop the application of the GMM to the analysis of single molecule dwell times. We present a general method that can be used to create objective functions applicable to single molecule reactions. We characterize the performance of our method by testing it on simulated data. We simulate several common reaction schemes, including single, double, and triple step reactions. We have characterized the bias and dispersion of the estimates generated by our method and compared these to the most common alternative method of analysis, NL-LSQM. Additionally, we have collected experimental data on site-specific DNA cleavage by the restriction endonuclease NdeI and analyzed the measured dwell times using our method.

## Theory

### Motivation

More detailed reviews of the GMM can be found in the literature (9). Here, we review the essentials necessary to understand our work. As motivation, we consider a system with two hidden parameters, *a* and *b*, that define a probability density *p(t;a,b*) for a random variable *t*. Define *m*_1_(*a,b*) = 〈*t*〉 and *m*_2_(*a,b*) = 〈(*t* − 〈*t*〉)^2^〉, where the angular brackets indicate the population mean (also known as the expectation value). The population mean of any function *f*(*t*) is here defined as

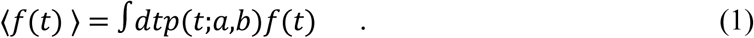

Note that in Eq. 1, the population mean has an implicit dependence on the parameters *a* and *b*. These two functions *m_1_*(*a*,*b*) and *m*_2_(*a*,*b*) are simply the population mean and variance of the random variable *t*. Now, let **t** be a T-dimensional vector whose elements are samples of the random variable *t*. In this paper, variables in italics represent real numbers and variables in bold represent vectors. The moment functions

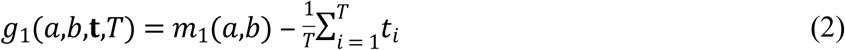

and

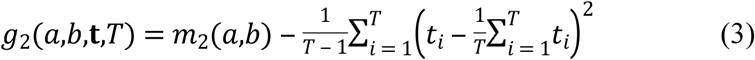

both express the difference between a population moment (first term on the right hand side) and a sample moment (second term on the right hand side). Notice that the unbiased estimator is used for the variance (Eq. 3). If sample moments give good estimates, then we expect the values on the right hand sides of Eqs. 2 and 3 to be very small for the true values of *a* and *b*. A reasonable method would be to set the right hand side equal to zero and solve for *a* and *b*. This method of generating *M* moment equations for *M* parameters (in this case *M* = 2) is the Classical Method of Moments (CMM). There are limitations. There may not be real solutions to the equations. Additionally, there may be multiple conditions, i.e., moment equations that should be satisfied. In this case, the number of equations may exceed the number of parameters (the over-constrained case), producing an overconstrained system of equations that may have no solution.

### General formulation of the GMM

The GMM is a general framework for solving the problems identified in the previous section. In the following derivation, we assume a single random variable. This case generalizes in a straightforward manner to include multi-variate random processes. In our derivation, we assume a probability density for the random variable. It is not strictly necessary that the functional form of the probability density is known. As we will show, our application can be used to determine the *moments* of the distribution without knowing the density. In practice, to relate the moments to questions such as “how many steps are needed to explain this data?” one needs a functional form of the density. Therefore, a single application of this method cannot by itself answer the question “which is the best model to describe this system?” However, comparison of analyses using different models can help with this important problem, as we will show later in this work.

To start, we assume the probability density is a function of the random variable *t* as well as of *N* parameters **λ** and will be written as *p*(*t*; **λ**). Note that **λ** has dimension *N*. We will also need the joint probability density for multiple independent samples. If we assume there are *T* samples, this is

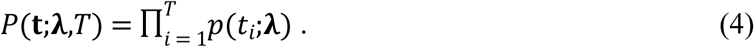

We also need the expectation value of a function of t. This is defined (analogously to Eq. 1) as

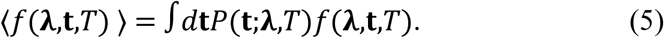

In Eq. 5, the possible dependence on **λ** and *T* has been explicitly included.

In order to determine a GMM estimate of the parameters from a given sample, we start with a set of functions, termed generalized moments, such that the expectation value of these functions is zero. That is, we need a set of *M* functions *g_m_*(**λ**,**t**,*T*) such that 〈*g_m_*(**λ**,**t**,*T*)〉 = 0 In practice, it is these functions we must assume, not the probability density. However, if we know the probability density, it is straightforward to determine the generalize moments. The index *m* can assume any value from 1 up to *M* (the maximum number of moments considered). In general, we may have more moment conditions than parameters (*M* > *N*), and therefore we can only look for approximate solutions. To do this, the following objective function is minimized with respect to the variables **λ**,

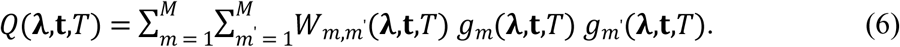

The matrix *W_m,m’_* is a positive definite weight matrix which in general can depend on the parameters **λ** as well as on **t** and *T*. The final GMM estimates of the parameters can then be written as

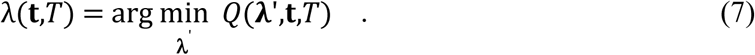

Note that the value of the estimate depends not only on the samples **t** and sample size *T*, but also on the choice of weight matrix *W_m,m’_*.

When the number of moment conditions *M* is less than the number of parameters *N*, this problem is underspecified, and the function *Q* does not have a unique minimum. If the number of moment conditions is equal to the number of parameters, the system is “just specified,” and, in general, the GMM will give identical estimates as the CMM. This latter point follows since *W_m,m’_* is positive definite and therefore *Q* ≥ 0. Since the CMM estimates make *g_m_* = 0 for all *m*, we have *Q* = 0 identically and therefore it must be a minimum at that point. It is important to remember that this theorem holds only when the CMM estimate exists (i.e., there are real solutions). Note that it also follows that the just-specified case will be *independent* of the weight matrix *W_m,m’_*, as the CMM does not use that matrix. In general, for the over-specified case, the GMM estimate does depend on choice of weight matrix, a problem to which we now turn.

In the limit of a large number of samples, the distribution of moments (for fixed parameters), is expected to be Gaussian as a consequence of the Central Limit Theorem. In this case, choosing a weight matrix equal to the inverse of the covariance matrix of the moment functions will lead (in the limit of a large number of samples) to an unbiased estimate of parameters with minimal variance (11). One should remember that, in general, for finite numbers of samples, the estimate will be biased and not normally-distributed.

To see how this is applied, define the following covariance matrix

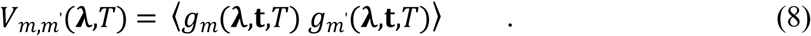

Here, as before, the angular brackets indicate the population mean. Note that since we integrate over the random variables, this covariance matrix is not a function of **t**. However, it may depend on sample size, *T*. The weight matrix is then the inverse of this matrix:

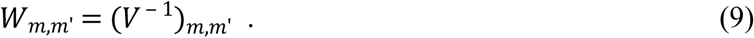

One problem with this approach is that the weight matrix is a function of the parameters **λ**. This complicates the minimization of the objective function, since the values of these parameters are unknown. Various practical methods of handling this include (1) using the identity matrix for *W_m,m’_* (a simple method for getting an initial solution), (2) using various estimates of the covariance matrix calculated from the data itself, (3) using multi-step methods where a simple weighting is used first (often the identity) and then the resulting GMM estimate for the parameters is used to recalculate the weights for a second pass, or (4) continuously updating the weight matrix as the objective function is minimized. With the exception of the continuously updated weight matrix, we will evaluate all of these methods in this work.

### Application to single molecule dwell times

The reactions considered will be assumed to be of the type shown in Fig 1. In this figure, states of the system are represented by letters, and the corresponding mean residence times in each state are given by τ_A_, τ_B_, etc. For the reversible reaction, transition rates are given as k_1_, k_2_ and k_3_. The observed dwell time is defined as the time it takes the system to enter the final state given that it starts out in state A. For the derivation that follows, we will assume all reactions steps are irreversible. Since the probability density of the total dwell time for a two step reversible reaction has the same functional form as that of an irreversible scheme, our derivation is applicable to both cases. We will show at the end of this section how to apply our analysis to the reversible two step reaction. For more than two steps, the functional form of the probability density for reversible schemes is more difficult to relate to our method. We choose to limit ourselves to the irreversible case for more than two steps for two main reasons. First, the theoretical form of the cumulants can be expressed generally and simply, thus allowing a simple and general formulation of a solution. Second, testing N-step irreversible reaction schemes against data is an established method for estimating the number of steps present in experimental data (12, 13), thus allowing for application of the cumulant based GMM method to address this question.

**Fig 1.**
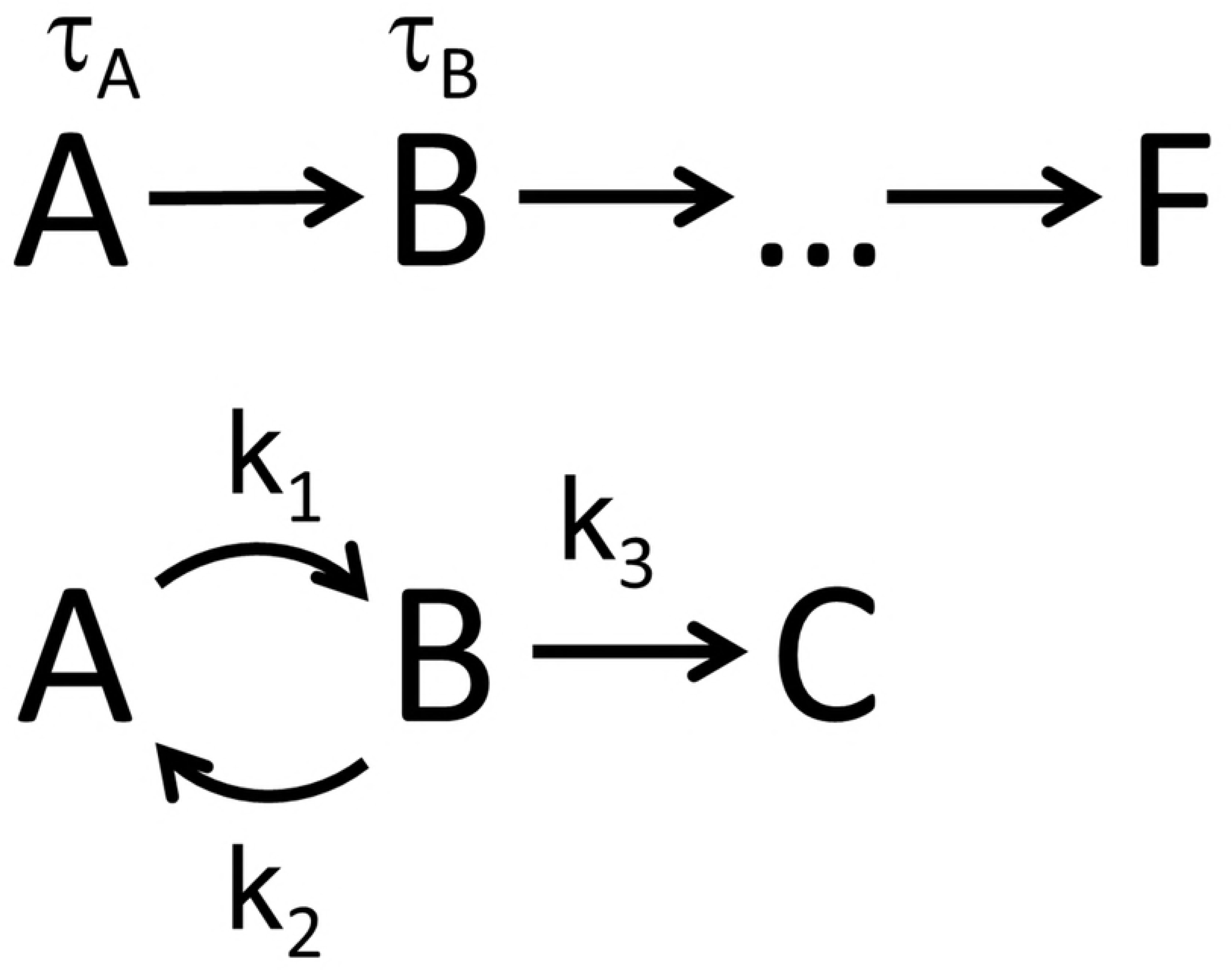
Reaction schemes. Each state is represented by a letter and the mean residence time in that state is τ_A_, τ_B_, etc. The transition rates in the reversible reaction are given by k_1_, k_2_ and k_3_.

The residence time t_A_ in state A in Fig 1 is drawn from a continuous, single-exponential distribution,

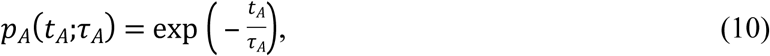

where τ_A_ is the mean dwell time in state A. A similar function can be defined for state B, C, etc. The mean observed dwell time for the system is therefore

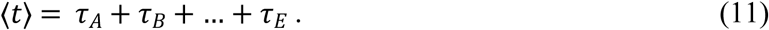

Since each step is independent and exponentially-distributed, the population variance is given by an analogous formula,

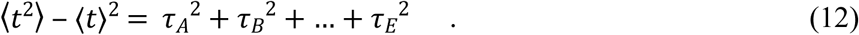

In order to formulate a general method for these systems, we would like a formula analogous to Eq. 12 for higher order moments. We can find such a generalization with cumulants. For any random process which is a sum of independent random processes, the *m*^th^ order cumulant is merely the sum of the *m*^th^ order cumulants of the individual processes. In this case, if we represent the cumulant of the total dwell time with κ^(m)^, then we have

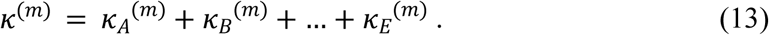

In this equation, κ_A_ ^(m)^ is the *m*^th^ order cumulant of step A, etc. The *m*^th^ order cumulant for an exponential distribution is

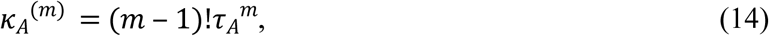

which allows us to write the general formula for the cumulants for an *N*-step irreversible reaction scheme.

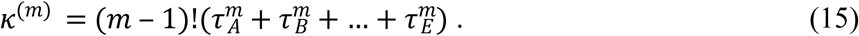

Eq. 15 gives the population cumulants of the reaction scheme shown in Fig 1. In order to complete the moment function, we need to calculate unbiased estimates of the cumulants from the measured samples. Formulas for unbiased sample cumulants exist and are referred to as *k*-statistics (14). In order to define the *k*-statistics for this problem, let us first define the *m*^th^ order central moments.

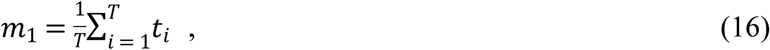

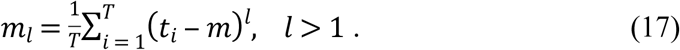

The *k*-statistics up to order 4 are then

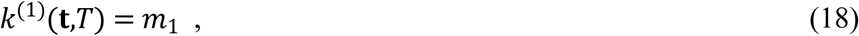

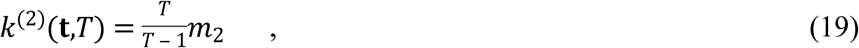

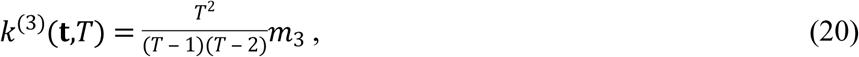

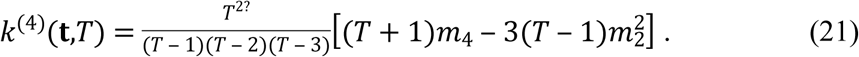

The first and second order expressions are the usual definitions of the sample mean and sample variance in the random variable *t*. The third and fourth order expressions are less familiar.

We can now state our *m*^th^ order generalized moment function for the *N*-step reaction.

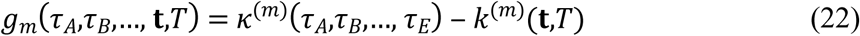

In this expression, the population cumulant *κ*^(*m*)^ is given by Eq. 15 above, and the term *k*^(*m*)^ is to be an unbiased estimate of the cumulant as calculated from Eqs. 18 – 21.

The GMM is not completely defined until the weight matrix is specified. We evaluate several options in this paper. These include the identity matrix, the inverse of a jackknife estimate of the covariance matrix, and the inverse of a covariance matrix calculated using a Monte-Carlo method. We leave the description of these methods to the Materials and Methods section of this paper. Equations 14, 15, and 16-21 completely define the moment functions used in this study. They, along with the weight matrix, define the GMM for this problem.

### The case of a reversible two step reaction

For a two-step reaction, such as those shown in Fig. 1, the total dwell time can be shown to obey a probability density of the following form (12),

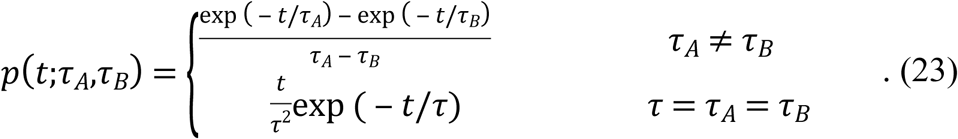

In the case of a reversible first step, the decay constants will be equal to

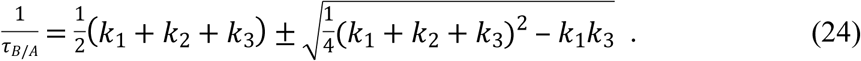

In this equation, the plus sign applies to τ_B_ and the minus sign to τ_A_. The GMM described in this work will determine estimates for the parameters τ_A_ and τ_B_, which can be related to the underlying rate constants in the reversible reaction model by Eq. 24.

## Materials and methods

### Simulations and GMM calculations

Dwell times were simulated for multistep stochastic reactions by adding samples individually drawn from exponentially-distributed random processes. The sets of decay parameters used were as follows. For the single step, all trials used a decay time of 10 s. For the two-step reaction, decay parameters pairs used were (10 s, 10 s), (10 s, 20 s), (10 s, 50 s) and (10 s, 100 s). For the three-step reaction, decay parameters sets were (10 s, 10 s, 10 s), (10 s, 10 s, 50 s), and (10 s, 30 s, 100 s). For all decay parameter sets, sample sizes used were 5, 10, 20, 50, 100, 200, 500 and 1000. For each set of parameters and sample size, 1,000 trials were simulated. For each trial, unbiased estimates of cumulants were calculated using *k*-statistics up to 4^th^ order.

Four different weight matrices were evaluated. These were (1) the identity matrix, (2) a diagonal matrix with elements equal to the inverse of a jackknife estimate of the variance of the cumulants, (3) the matrix inverse of a jackknife estimate of the covariance matrix of the cumulants, and (4) the inverse of an interpolated covariance matrix. All jackknife estimates were determined from a single trial in the following manner. First, N subtrials were created by sequentially removing each sample from the trial (resulting in N subtrials of N-1 samples each). Then, these N subtrials were used to calculate N distinct estimates of the cumulants. Finally, the variance or covariance was calculated from these estimated cumulants and the result was scaled by (N-1)/N. The interpolated covariance was calculated by linear interpolation from a set of previously-calculated covariance matrices. These previously-determined matrices were calculated using a Monte Carlo (MC) method for a wide range of values of N (sample size) and decay parameters (τ_A_, τ_B_, etc.). For the MC calculations, 100,000 trials were calculated, and the covariance was calculated directly from the cumulants of these trials.

The objective function (Eq. 6) was minimized using the Broyden-Fletcher-Goldfarb-Shanno algorithm with an explicit gradient. Global minima were found by using a logarithmically-spaced grid of initial trial points. The parameter space region searched was an ‘n-cube,’ where n is the number of parameters (the number of steps in the reaction). The grid spacing was a factor of ten. For example, for a two-step reaction, the two dimensional region [1, 1000] × [1, 1000] was searched. This included starting points an order of magnitude less and more than the minimum and maximum decay parameters simulated. Note that since the GMM objective functions examined here are multipolynomials of the decay parameters, the minimization is robust, as the surface is not rough (i.e. it does not exhibit a large number of minima). In the two-pass GMM, the estimates for the decay parameters from a first-pass minimization were used as the starting search point for a second pass which used the interpolated weight matrix calculated from the first pass parameter estimates.

All scripts and routines for performing the calculations reported in this manuscript can be downloaded from https://sourceforge.net/projects/genmm. These scripts are not intended to be end user software, but are made available to encourage transparency, reproducibility and to encourage others to build on our work.

### Nonlinear least-squares minimization

To perform the NL-LSQM of the simulated data, each simulated data set was binned into a number of bins equal to the square root of the number of samples (rounding up to the nearest whole number). The bins were fit to a bi-exponential distribution with two free decay parameters using the Levenberg-Marquardt algorithm. A finite bin-width correction was used.

### Experimental data collection and processing

Experimental dwell times were measured for DNA cleavage by the restriction endonuclease NdeI. The method of data collection is explained in detail elsewhere (6). Briefly, 1000 bp DNAs with a single centrally-located NdeI restriction site were used to tether 1 μm magnetic beads in a microflow channel. The DNA-tethered beads were observed under low magnification and dark field imaging. A low flow rate and magnet were used to apply a small force (<100 fN) to the beads, which are pulled out of focus as the DNA is cleaved. Varying concentrations of NdeI (from 25 to 350 pM) were introduced in reaction buffer (20 mM Tris-HCl, 100 mM NaCl, 3 mg/mL BSA, 1 mg/mL Pluronic F-127, 1 mM MgCl_2_). Video data was recorded at a frame rate of 1 fps. Dwell time between initial introduction of the enzyme and the final cleavage of the DNA is measured by noting the time at which the bead is removed. The software package ImageJ was used to locate bead positions in the initial image, and then all images were analyzed by custom software which integrates the intensity around the bead position. A large drop in intensity identifies the cleavage event and the dwell time is recorded as the frame number. A frame rate of either 0.5 or 1 fps was used. Individual data sets produced 200 to 700 dwell times (cleavage events). Mean dwell times varied from 150 s to 300 s, and data was analyzed using the GMM. One up to six step reaction models were tested using the “just-specified” objective function for each scheme. A single pass GMM was used.

## Results

### Performance of GMM with simulated data

By minimizing an objective function composed of moment functions, the GMM provides estimates of parameters which approximately satisfy the condition that all the moment functions have the value zero. The performance of the GMM is known to depend on the order of the method (the number of moment conditions included), as well as on the nature of the weighting matrix used to weight the various terms. In addition, various methods of weighting have been developed. These include “two-pass” methods, in which an initial trail matrix is used to find a first pass estimate. The second pass uses a more effective weight matrix calculated from the results of the first pass.

In this work, we apply an algorithm based on the GMM to the analysis of single molecule dwell times. We use simulated as well as experimental data, and test a variety of models, weighting matrices, as well as a two-pass method. See the Materials and Methods section for a complete description of all variations. In general, we found a diagonal weight matrix based on jackknife estimates of the variances in the cumulants (the D-matrix) to produce estimates with the lowest bias for all orders. Figure 2 shows the results of analyzing simulated data for a single step reaction. The results are shown for methods of all orders up to fourth. The dispersion in the results is shown by representing the mean deviation with error bars. The unit matrix (I-matrix) as well as a non-diagonal matrix based on jackknife estimate of the full covariance matrix of the cumulants (C-matrix) gave poorer results (data not shown).

**Fig 2.**
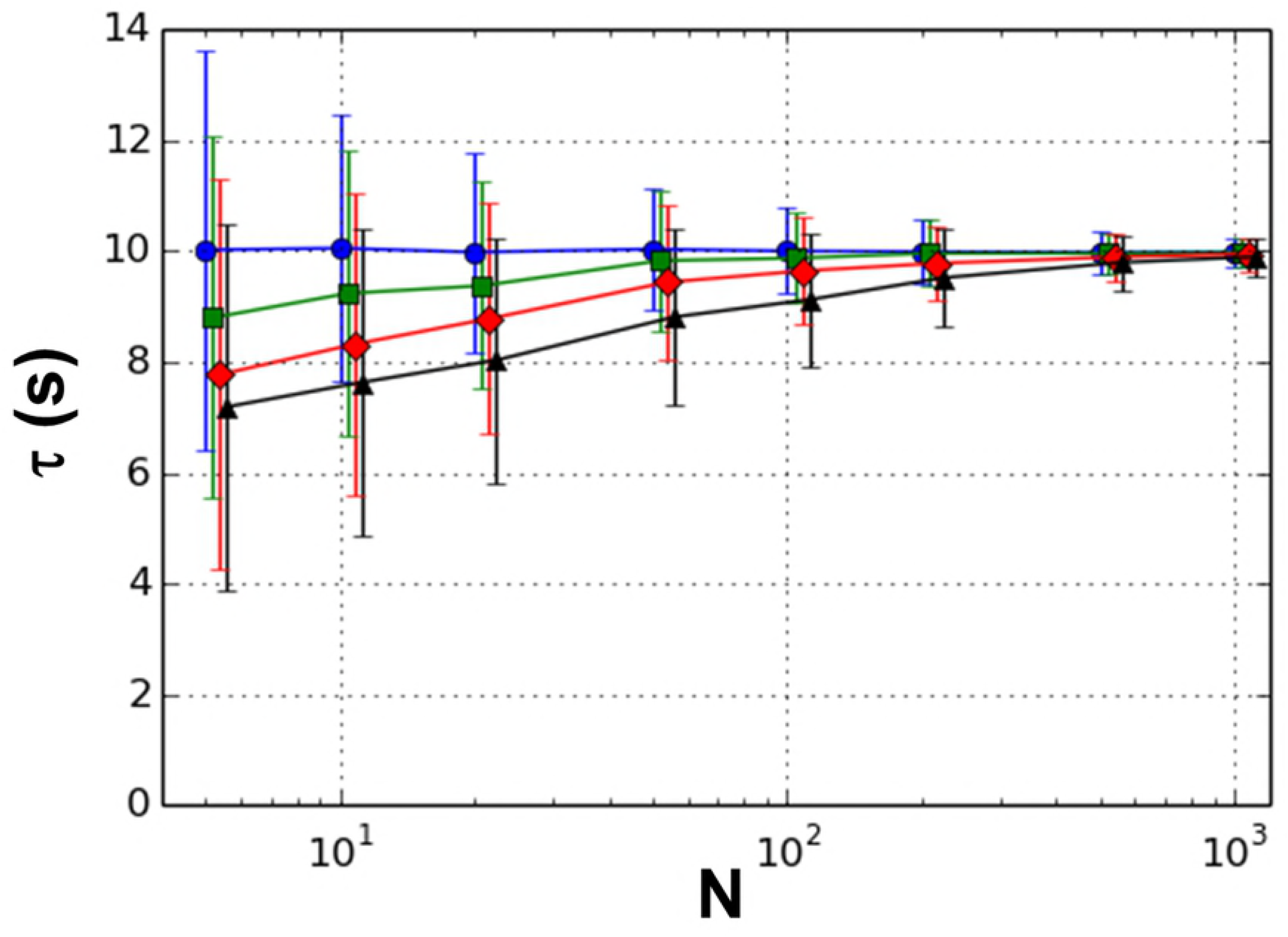
Results for 1 step reaction. Mean for 1000 trials for orders 1 (circles), 2 (squares), 3 (diamonds), and 4 (triangles) are plotted versus sample size (N). The simulated decay constant is 10 s. Error bars represent mean deviations.

As seen in Fig 2, the bias and dispersion reduce with increasing number of samples for all orders (except for first order where the bias is zero). Additionally, the bias tends to increase as the order of the method is increased for all sample sizes. Note that the first order method is equivalent to the Classical Method of Moments, in which the estimate is simply the mean of the sample dwell times. It is easily shown that this estimate is unbiased for all sample sizes. We found that the dispersion, as measured by the mean deviation, shows a complex dependence on the order of method for the I-matrix. However, for the D- and C-matrix, the dispersion is relatively independent of order (data not shown).

We now turn to the performance of the GMM for the two-step reaction (a single intermediate). In all that follows, we hold the first decay parameter fixed (τ_A_ = 10 s) and vary the second (τ_B_ = 10, 20, 50, or 100 s). We examined second, third and fourth order methods. Note that in this case, the first order method is under-constrained and does not result in a unique estimate and is therefore excluded. We also examined the effect of using the I-, D-, and C-matrices (defined above), as well as the benefit of a two-pass GMM, in which a first pass estimate is used to recalculate the weight matrix.

For the two step reaction model, the GMM returns two estimated decay parameters which can be sorted into a smaller value (presumably an estimate for τ_A_) and a larger value (presumably an estimate for τ_B_). In Fig 3, we plot the mean values of these two estimates, along with the mean deviations as error bars, for two sets of model parameters, (τ_A_, τ_B_) = (10 s, 10 s) and (τ_A_, τ_B_) = (10 s, 50 s). The results shown were calculated using the D-matrix, which gave the best results for these trials (see Figs S2 through S6, Supplementary Material, for C and I-matrices). The bias and dispersion decrease with increasing sample size, consistent with the expected large sample size behavior (4). For the case of τ_B_ = 50 s (Figs 3C and 3D), we see that for both estimates, bias *increases* with order, similar to the pattern for the one step reaction. However, the sign of the bias is different for the two estimates, such that the lower estimate is too high and the higher estimate is too low. Dependence of the dispersion on the order is either weak or shows a slight increase as order goes up. Results for decay parameters τ_B_ = 100 s are similar to those for τ_B_ = 50 s (see Figs S1 through S6).

**Fig 3.**
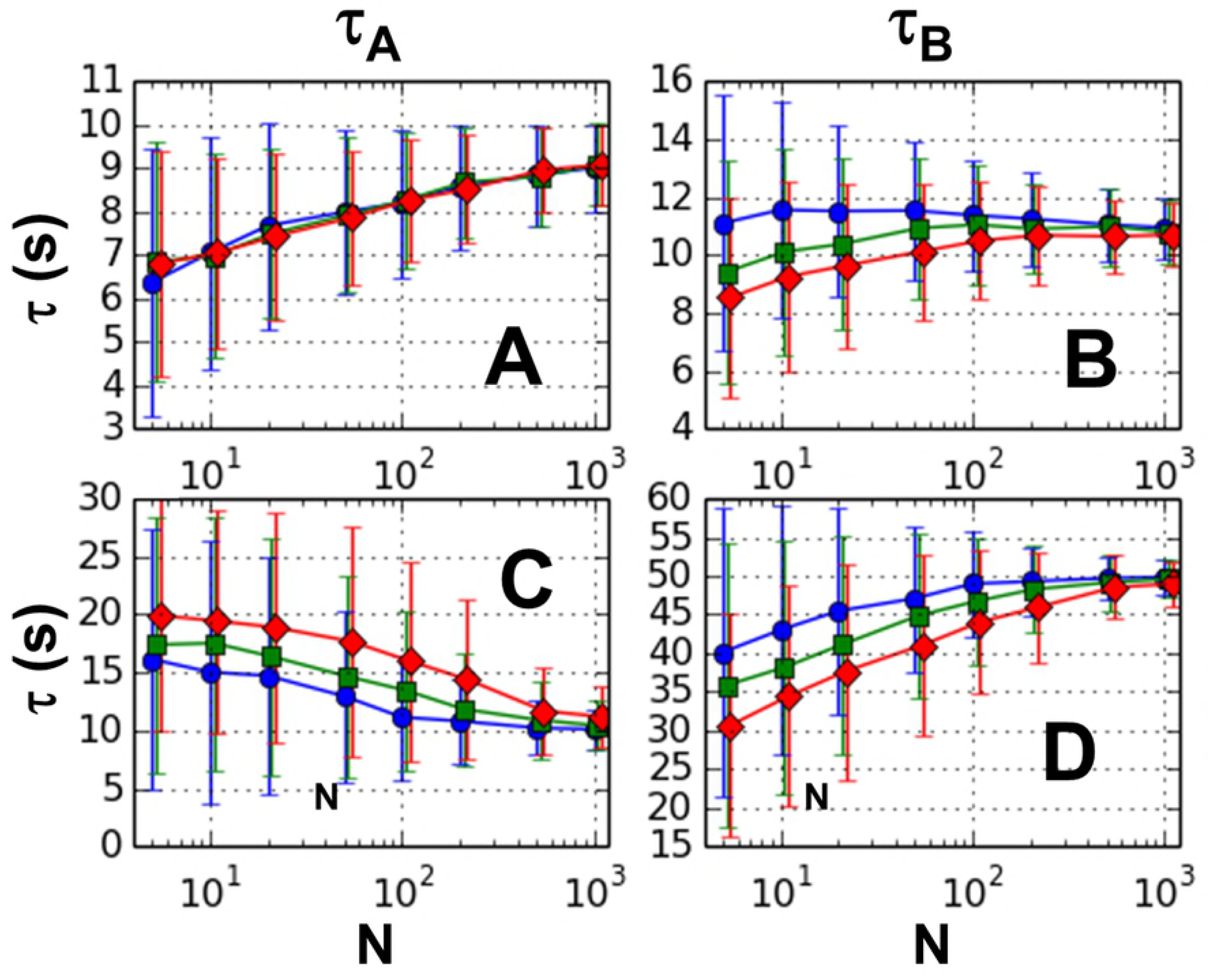
Single pass GMM results for 2 step reaction. Means for 1000 trials for orders 2 (circles), 3 (squares), and 4 (diamonds) are plotted versus sample size (N). Error bars represent mean deviations. Left panels (A and C) show lower estimate and right panels (B and D) show higher estimate. The two top panels (A and B) are for the decay constant pair (10 s, 10 s) and the two bottom panels (C and D) for (10 s, 50 s).

For the case when both model parameters are equal (τ_A_ = τ_B_ = 10s), the bias is either independent of order (Fig 3A), or shows a non-trivial dependence on order and sample size (Fig 3B). Results for decay parameters (τ_A,_ τ_B_)= (10s, 20s) are intermediate between those for τ_B_ = 10s and τ_B_ = 50s (see Fig S1). The above results indicate that a second-order method using the D-matrix is the optimum method of those explored using the parameters we tested.

We next tested the two-pass GMM on the simulated data. Two-pass GMM methods attempt to generate a more accurate weight matrix by executing a first pass using a best guess for the weight matrix, and then using the resulting first pass estimates to calculate a more accurate weight matrix for the second pass. In our case, we chose our best single-pass GMM result (the second order, D-matrix method) as our first pass. We then used the estimates from this pass to calculate the theoretical covariance in the sample cumulants and from the inverse of this, the weight matrix. For efficiency, the theoretical covariance was calculated by interpolation from a set of covariance matrices that were calculated using a Monte Carlo method (see Materials and Methods for details). Since the second order GMM is “just-specified” in the case of two model parameters, and hence independent of weight matrix, we investigated the effect of adding a second pass using up to third or fourth order moments

The results from the two-pass GMM are shown in the Fig 4. As can be seen from the figure, an additional pass does not greatly affect the bias nor the dispersion under any of the parameter sets used in this study. The small reduction in bias seen at low sample numbers in Figs. 4A and 4D is offset by the small increase in bias shown in Figs. 4B and 4C. Figure 4 only shows the results from the D-matrix weighting scheme. See Supplementary Material (Fig S7) for complete results.

**Fig 4.**
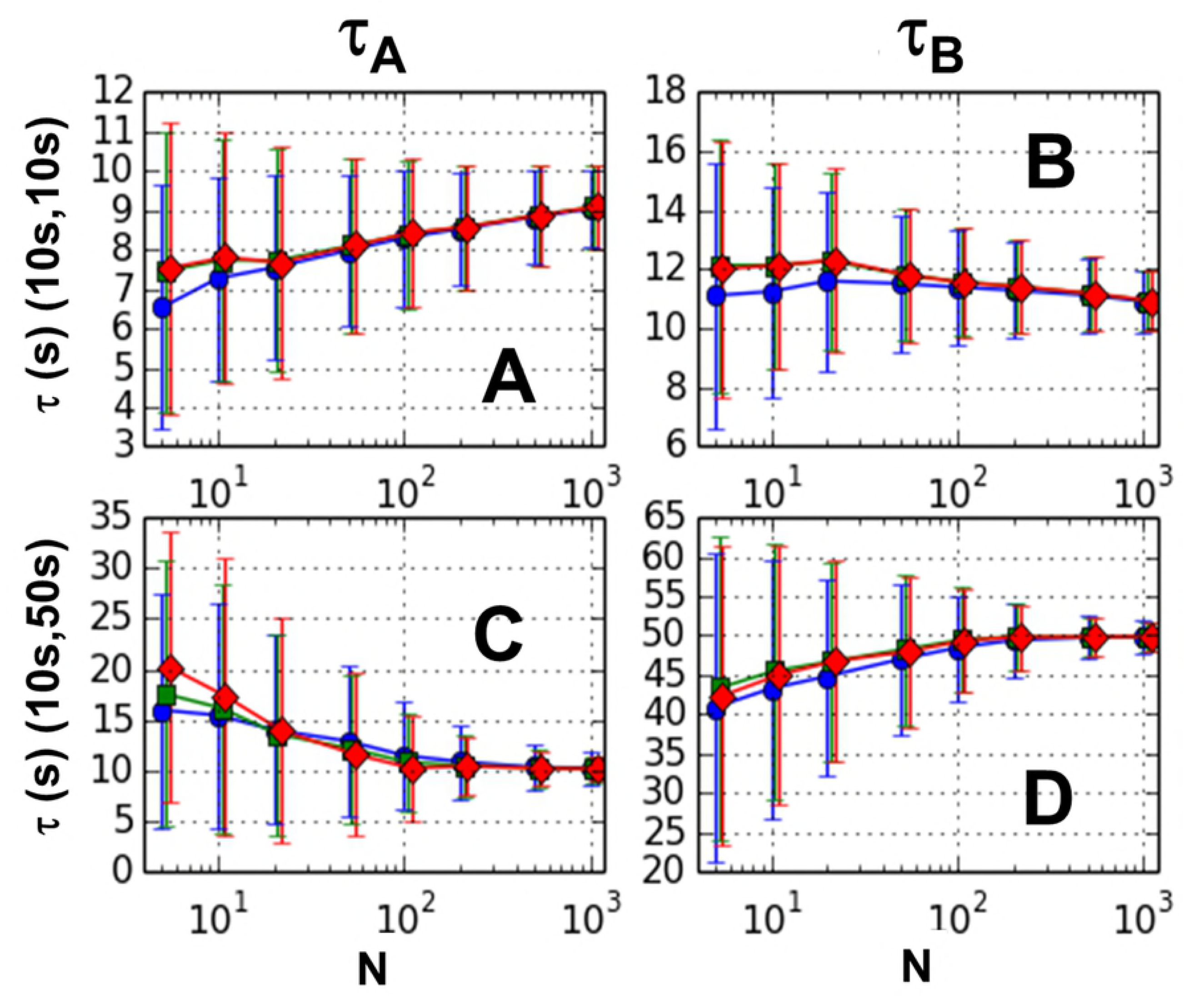
Two pass GMM results for 2 step reaction. Mean for 1000 trials for one-pass 2^nd^ order (circles), two-pass 3^rd^ order (squares), and two-pass 4^th^ order (diamonds) is plotted. Error bars represent mean deviations. Left panels (A and C) show lower estimate and right panels (B and D) show higher estimate. The two top panels (A and B) are for the decay constant pair (10 s, 10 s) and the two bottom panels (C and D) for (10 s, 50 s).

We additionally applied the GMM to a three-step reaction model. Third and fourth order single-pass GMM methods were applied to three sets of decay parameters. These sets were (τ_A_, τ_B_, τ_C_) = (10 s, 10 s, 10 s), (10 s, 10 s, 50 s) and (10 s, 30 s, 100 s). GMM estimates were sorted into smallest to largest returned value, and the means and mean deviations are shown in Fig 5. The performance of the GMM estimation is quite varied when faced with this more challenging problem. Bias tends to decrease with increasing sample number for most estimates, except for the lowest estimate for the case of parameters (10s, 10s, 50s).

**Fig 5.**
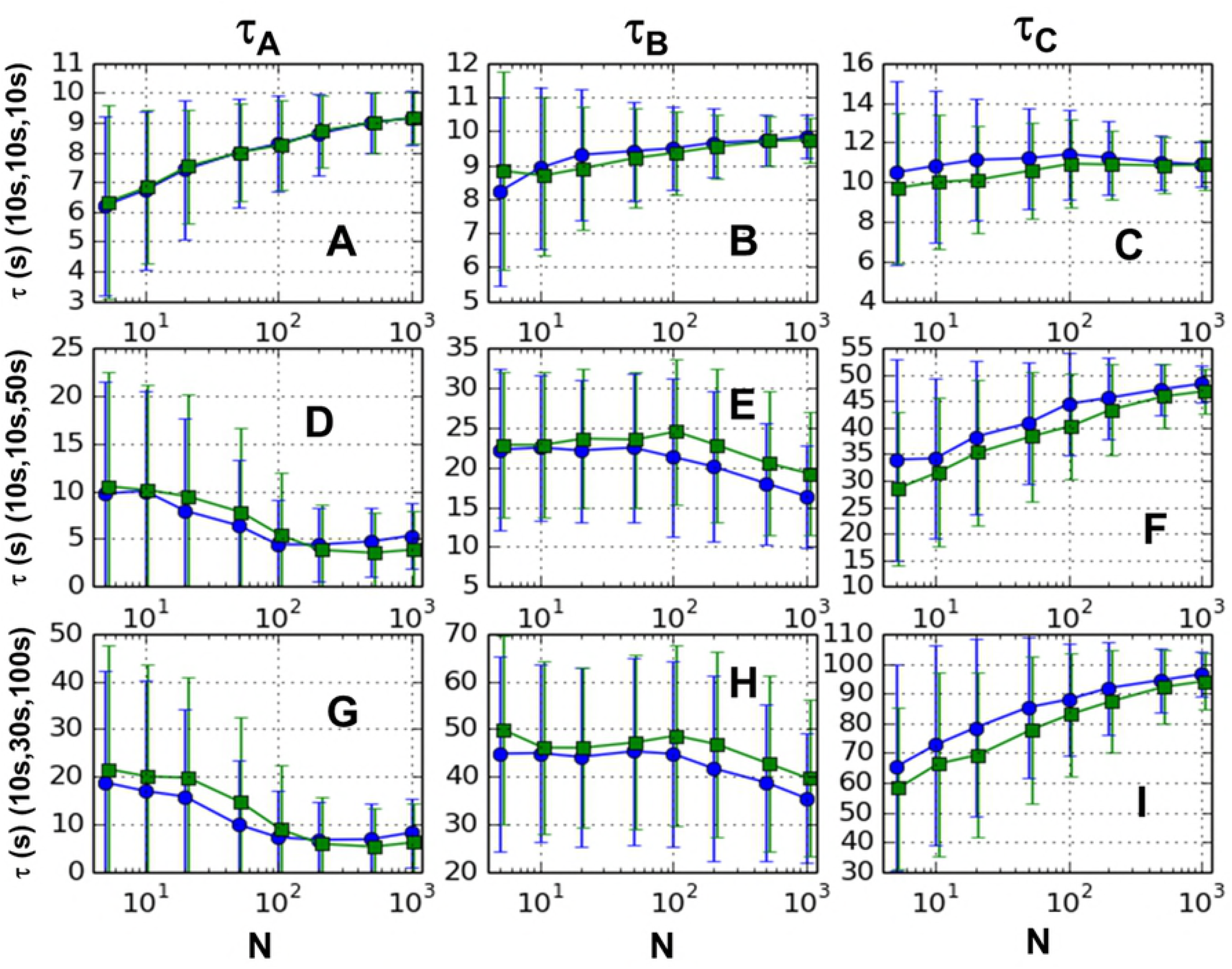
Single pass GMM results for 3 step reaction. Mean for 1000 trials for orders 3 (circles) and 4 (squares) versus sample size (N). Error bars represent mean deviations. The three GMM estimates are sorted into smallest to largest and arranged left to right in each row. Top panels (A, B, & C) are for the decay constants (10s, 10s, 10s), middle panels (D, E, & F) for (10s, 10s, 50s,) and bottom panels (G, H, & I) for (10s, 30s, 100s).

### Comparison with NL-LSQM

In order to compare the GMM to alternative methods, we used a non-linear least squares minimization (NL-LSQM) method based on fitting histograms of our simulated two-step reaction data to the theoretical bi-exponential distribution. We used the same simulated data that was used to calculate the GMM estimates shown in Fig 3 and performed global nonlinear least-squares minimizations as described in the Materials and Methods. Figure 6 shows a comparison of the second order GMM method using the D-matrix to the NL-LSQM method for a two-step reaction. Note that for sample sizes of 5 and 10, the NL-LSQM method showed very large bias for the smaller decay parameter (in some cases, returning negative decay parameters) which is not plotted. For large samples (N > 100) the two methods are comparable in bias and dispersion, except for the case where the two model parameters are equal, in which case the GMM has slightly smaller bias. At low sample size (N < 20), the GMM shows less bias and dispersion, and is able to return reasonable estimates even for sample sizes as small as five. Results for the decay parameter pairs (10s,20s) and (10s,100s) are shown in Fig S8 in Supplementary Material.

**Fig 6.**
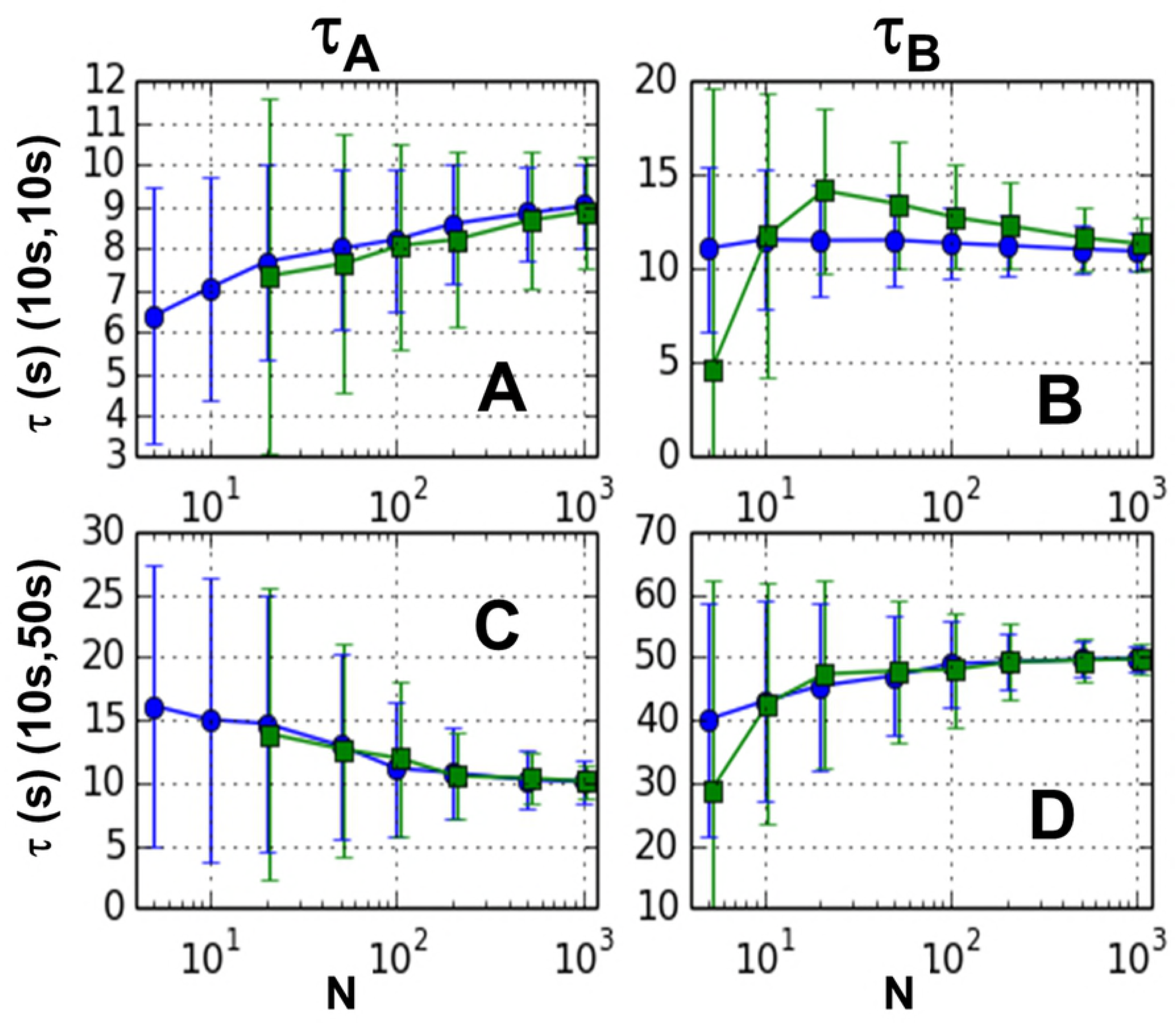
Results for comparison of GMM to non-linear least squares. Estimates from GMM are shown in blue circles and those of NL-LSQM are green squares and are plotted versus sample size (N). Error bars represent mean deviations. Left panels (A and C) show lower estimate and right panels (B and D) show higher estimate. The two top panels (A and B) are for the decay constant pair (10s,10s) and the two bottom panels (C and D) for (10s, 50s).

### Application to experimental data

To gain experience using the GMM with experimental data, we collected single molecule data of double stranded DNA cleavage using a bead loss assay we have previously described (6). Single molecule dwell times of DNA cleavage have been shown to be useful for the study of restriction endonuclease mechanism (15). Our technique uses tethered beads to measure the dwell time until cleavage for several hundred DNAs in a single experiment. The total dwell time consists of several steps, including the DNA target search as well as the cleavage of each strand of the DNA.

We have previously shown that the mean dwell time under the conditions of 2 mM Mg^2+^ is highly dependent on protein concentration, consistent with a diffusion-controlled process. In this work, we collected data at 1 mM Mg^2+^ for a range of concentrations from 25 pM to 350 pM and analyzed the resulting dwell times using the GMM. Since we did not know *a priori* the number of steps in the reaction, we chose to test models with one up to six steps, each using the “just specified” order (i.e., first order for one step, second order for two step, etc.) and using a diagonal jackknife weight matrix.

Our results for the one and two step analysis are shown in Fig 7. The single step result is equal to the sum of the two decay times from the two step analysis, which must be the case. The results for the three and four step method are listed in Table I. The results for the five and six step model are not shown. Examination of the graph and table shows that the slowest time step remains relatively constant for the different models as we increase the number of steps in the model past two. For example, at 22 pM, the slowest step in the two step model is 242 s, which changes to 259 s in the three step model, then 265 s in the four step reaction. But the faster steps show a different pattern. For this same data, the two step model yields a faster step of 72.0 s, which turns into two steps of 27.5 s in the three step reaction, and finally three steps of 16.4 s in the four step. Note that in each model, the faster steps are each of identical duration. This pattern of the slowest step remaining somewhat constant and the faster steps all being equal in value, but decreasing as we increase the number of steps in the model continues for the five and six step models also. This is a curious finding, and suggests that this pattern might help distinguish how many steps are the minimum necessary to explain our data.

**Table 1.** Results of three and four step GMM analysis of experimental data.

| [NdeI] | THREE STEP RESULTS |  |  | FOUR STEP RESULTS |  |  |  |
| --- | --- | --- | --- | --- | --- | --- | --- |
| (pM) | $\tau_1$ (s) | $\tau_2$ (s) | $\tau_3$ (s) | $\tau_1$ (s) | $\tau_2$ (s) | $\tau_3$ (s) | $\tau_4$ (s) |
| 22 | 27.5 | 27.5 | 259 | 16.4 | 16.4 | 16.4 | 265 |
| 44 | 11.1 | 11.1 | 199 | 5.05 | 5.05 | 5.05 | 206 |
| 88 | 21.4 | 21.4 | 145 | 12.4 | 12.4 | 12.4 | 151 |
| 175 | 11.8 | 11.8 | 159 | 5.96 | 5.96 | 5.96 | 164 |
| 350 | 36.3 | 36.3 | 91.9 | 21.6 | 21.6 | 21.6 | 99.7 |

**Fig 7.**
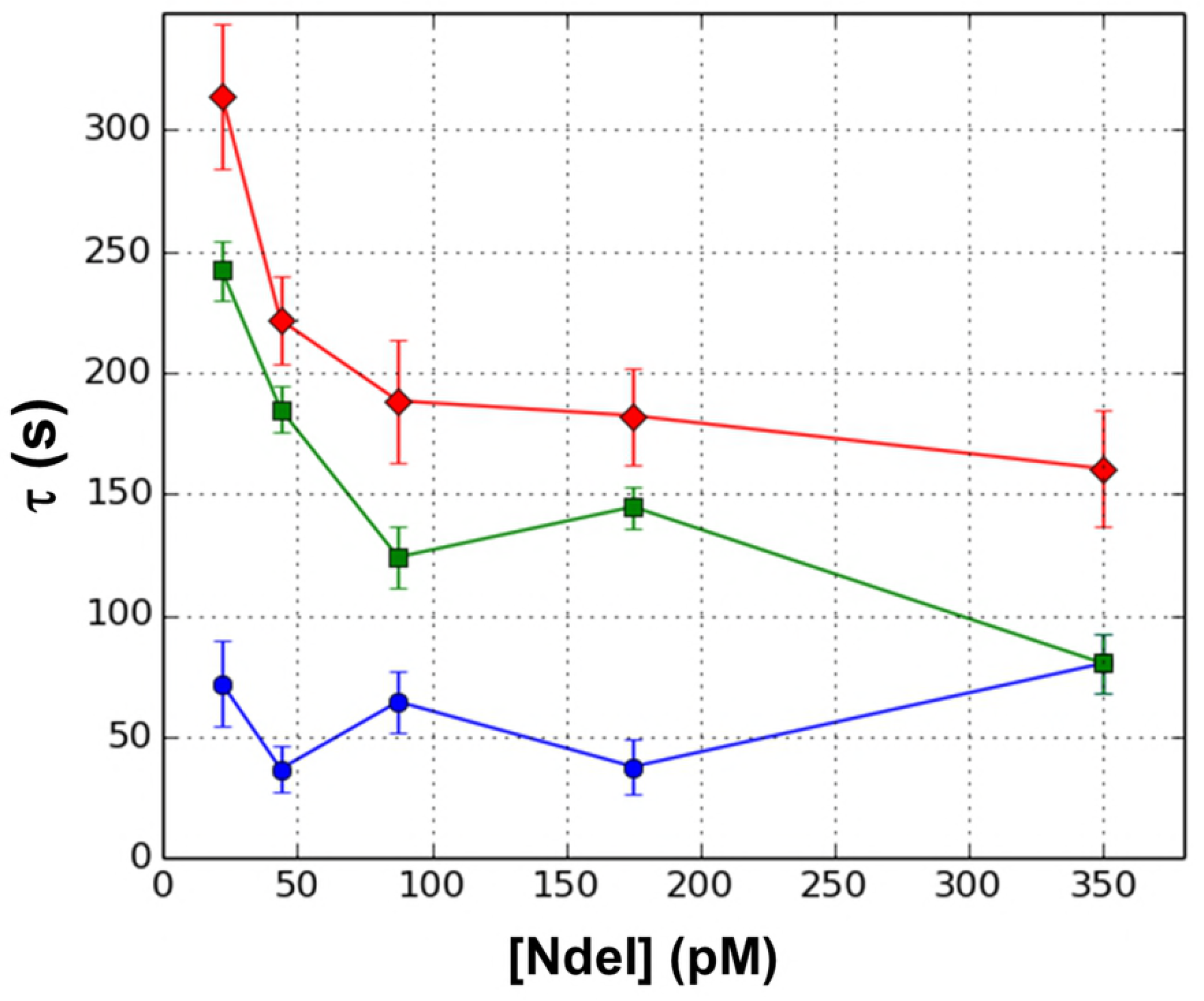
Application of GMM to DNA cleavage by NdeI. All results are for single-pass GMM. The results for 1 step reaction, first order GMM are shown with red diamonds. Results for a 2 step reaction, second order method are shown in green squares and blue circles. Error bars are estimates of statistical uncertainty taken from dispersion shown in Fig 3.

To explore this idea, we generated two simulated test datasets, one of a two-step reaction with time steps (50s, 150s), and one of a six-step reaction with time steps (10s, 10s, 10s, 10s, 10s, 150s). We then applied the just specified GMM (single pass) using different numbers of steps (one up to six) in the model. For the two step data, we found that as we increase the number of steps in the model, the method returned negative time constants when asked for more than two steps. The greatest returned decay times were relatively constant and close to (50s, 150s), indicating that even when we choose to analyze the data with the incorrect model, the method was correctly identifying the steps present in the data, and then returning insignificant durations for the fictitious steps. For the six-step simulated data, the largest time constant remained relatively constant and close to the expected value of 150s. The faster decay constants were all much smaller but varied quite a bit, and the algorithm began to return negative time constants when more than four steps were assumed in the model. This shows that the method could not correctly identify the number of fast steps, but did seem to suggest that there were at least four. This is similar to what we observed in the analysis of our data, and suggests that in our experimental system, there are several fast steps and one slower one. This slow step is shown by the the green curve in Fig 7 and the faster decays are represented by the blue curve in Fig 7, whose value is the sum of the faster decays.

## Discussion

We have developed an application of the Generalized Method of Moments (GMM) to the analysis of single molecule dwell times. Originally developed for the analysis of econometric data, the GMM is a statistical framework for the analysis of samples drawn from random processes. Our method is based on the analysis of cumulants of the data, and we have shown that it can be used successfully to analyze single molecule biophysical data. Using simulated as well as experimental data, we have shown that the GMM can extract useful model parameters directly from sets of experimentally measured dwell times. We also provide guidelines on the best use of the GMM to analyze real data.

Many of our findings agree with the growing literature assessing the utility of the GMM. Our simulated data shows that for applications to single molecule dwell times, the lowest order method that can completely determine a solution (the “just-specified” case) demonstrates lower bias than higher order methods. For orders higher than this minimum, the bias generally depends on the nature of the weight matrix. In the two-step reaction, we investigated the effect of three different weight matrices for these higher order GMMs. These were (1) the identity matrix, (2) a diagonal weight matrix with terms equal to the inverses of estimated variances of the cumulants, and (3) a matrix equal to the inverse of an estimated covariance matrix. We find that, although estimating the variances of the cumulants from the data can reduce bias, adding more complicated weight matrices (including off-diagonal terms due to cross-variances) can actually *increase* bias. These results indicate a “less-is-better” approach. Adding higher orders or off-diagonal elements to the weight matrix does not necessarily help the estimate. We also found that adding a second pass (see Fig 4) does not significantly improve performance.

Even under “just-specified” constraints, i.e., when the number of moment conditions equals the number of free parameters, our results for the two-step reaction show dependence on weight matrix, albeit small (See Fig S4 in Supplementary Materials). Although it is true that the just-specified GMM method generally gives the same results as the Classical Method of Moments (CMM), this only follows if the CMM returns a real-valued result. In the case of the two-step reaction, it is straightforward to show that the CMM fails to return real-valued estimates in many cases. In these cases, the GMM will return real values that *depend* on the weight matrix. Furthermore, it can be shown that in the cases in which the CMM does not return real valued estimates, the second order GMM must return a *double root*, that is, it returns two identical estimates. Since these estimates tend to be midway between the higher and lower decay constants, they have the effect of causing the higher estimate to have *negative* bias, and the lower estimate to have *positive* bias, a trend which is seen in our simulated data (Figs 3 and 4, lower panels). It is worth noting that other methods of dwell time analysis, including Bayesian methods and Hidden Markov Models, have been shown to systematically underestimate decay constants, similar to what we find for the GMM (7, 8).

Comparison of the GMM to NL-LSQM shows that these two methods provide estimates with comparable bias and dispersion for larger sample sizes (N > 20). However, the GMM shows a moderate advantage in terms of bias at very low sample numbers, and does better at estimating the *smaller* of two decay parameters (Figs 6A and 6C) in these cases. The NL-LSQM method often fails to pick out these faster decays when there are very few samples. A partial explanation can be found in the fact that the NL-LSQM method relies on binning which can be particularly challenging at low counts.

In our test of the GMM on experimental data, we found that as we added more steps to the reaction model, the method continued to produce estimates of faster multiple steps, out to six steps. This is not what we found for similar analyses of simulated two-step data, and suggests that in the experimental system, there are a number of faster steps but that the GMM is not able to determine the exact number of steps nor the rate of each one. Figure 7 shows that the slow step is dependent on the protein concentration and decreases as the concentration increases. However, the faster steps shows little dependence on protein concentration. This could be explained by a mechanism in which the rate limiting step is the site specific association of the protein with the binding site, followed by a series of faster steps leading to cleavage of one or both DNA strands.

Once implemented, the GMM is easy to apply with few adjustable parameters. It also is flexible and can be reformulated, requiring only a redefinition of the moment functions. The only requirement is that a sufficient number of moment functions of the measured values and system parameters can be formulated, whose expectation values are zero. The number of such moment functions must be at least the number of free parameters in the model.

## Acknowledgements

The authors would like to thank Stephen Gambino for his assistance in collecting the experimental data used in this work. This work was funded by the National Science Foundation RUI Award (MCB-1715317) and by Emmanuel College.

